# µCeta: a set of cetacean-specific primers for environmental DNA metabarcoding with minimal amplification of non-target vertebrates

**DOI:** 10.1101/2025.03.19.644246

**Authors:** Masayuki Ushio, Sachie Ozawa, Shin-ichiro Oka, Tetsuya Sado, Robinson O. Kisero, Lindsay Porter, Eszter Matrai, Masaki Miya

## Abstract

Biodiversity monitoring is crucial for understanding ecosystem dynamics and species distributions, particularly in the context of anthropogenic impacts and climate change. Cetaceans, as key indicator species of marine ecosystems, face increasing threats from human activities, highlighting the need for effective, non-invasive monitoring methods. In the present study, we developed novel Cetacea-specific primer sets to enhance the detection efficiency of cetacean species through environmental DNA (eDNA) metabarcoding, while minimizing the amplification of non-target vertebrates, such as fish and humans. We retrieved mitochondrial genomes of 71 cetacean species from a public database and designed 20 candidate primer sets, which were assessed *in silico* for their specificity and capacity to differentiate cetacean sequences. Four primer sets with the best *in silico* performance were selected for empirical validation using DNA from tissue samples and eDNA from seawater collected from aquarium pools and Hong Kong coastal waters. All four primer sets effectively amplified cetacean DNA from tissue samples. However, in the aquarium pool tests, three primer sets failed to accurately identify one or more cetacean species due to a lack of interspecific variation within the amplified region. From these, we selected one primer set targeting a 267 bp region of the mitochondrial 12S rRNA gene, named µCeta, and applied it to water samples collected from Hong Kong coastal waters, where the iconic Indo-Pacific humpback dolphin (*Sousa chinensis*) was observed. µCeta successfully detected *S. chinensis* eDNA while avoiding amplification of non-target species such as fish or humans. Our results demonstrate that µCeta is a reliable tool for cetacean eDNA detection in Hong Kong waters, contributing to cetacean conservation and enhancing our understanding of marine biodiversity.

## 1. Introduction

Biodiversity monitoring is crucial for conservation and ecological research, offering critical insights into ecosystem status, species distributions, and the impacts of ongoing human activities and climate change on ecosystem functioning (Cardinale et al., 2012). Ecosystem functions—such as biomass production, nutrient cycling, and sustainable food production— are directly linked to biodiversity (e.g., Hooper et al., 2005; Mace et al., 2012), underscoring the importance of maintaining diverse biological communities. Traditional monitoring methods, including direct visual censuses and camera and video traps, have long been used to assess species presence and abundance (Masuda et al., 2010; Nakagawa, 2019; Samejima et al., 2012). These methods provide invaluable data for enhancing our understanding, conservation, and management of ecosystems under increasing anthropogenic pressures. However, they are often labor-intensive and costly, requiring taxa-specific expertise, which limits their scalability and the monitoring frequency of biodiversity. Consequently, there is an increasing demand for complementary tools that provide more efficient and cost-effective approaches for biodiversity monitoring.

Environmental DNA (eDNA), defined as extra-organismal DNA directly extracted from environmental samples, has been utilized to detect the presence of macro-organisms (Ficetola et al., 2008; Miya et al., 2015; Taberlet et al., 2012, 2018). In macro-organisms, eDNA originates from various sources, including metabolic waste and damaged tissue (Kelly et al., 2014), and exists in different physical forms in the environment (Power et al., 2023). eDNA carries species-specific genetic information from the organisms that produced it, and eDNA analysis, such as metabarcoding, which utilizes universal primers and a high-throughput sequencer to obtain genetic marker sequences, enables detection of multiple species from a single environmental sample. This approach provides a non-invasive, cost-effective, and highly sensitive method for biodiversity monitoring in both aquatic and terrestrial environments (Bista et al., 2017; Bohmann & Lynggaard, 2022; Lynggaard et al., 2022; Ushio et al., 2017; Yamamoto et al., 2017).

Cetaceans, a term that includes all species of whales, dolphins, and porpoises, are iconic megafauna that inhabit all marine environments and serve as important indicators of the status of ocean ecosystems (Plön et al., 2024). However, many cetacean species are increasingly threatened by human activities and climate change (Lettrich et al., 2023; Li et al., 2023; van Weelden et al., 2021; Weir & Pierce, 2013). Their conservation is crucial not only for protecting marine biodiversity, but also for maintaining marine ecosystem functions and services, as they play critical roles in carbon sequestration, nutrient cycling, trophic cascades and eco-tourism (Estes et al., 2011; Kiszka et al., 2022; Roman & McCarthy, 2010). For instance, large, deep-diving cetaceans, such as baleen and sperm whales, contribute to vertical carbon and nitrogen cycling through their foraging behavior (Pearson et al., 2023). Smaller cetaceans also contribute to both horizontal (e.g., from coastal areas to the open ocean) and vertical transport of carbon and nutrients (Kiszka et al., 2022). Given their ecological significance and vulnerability (more than a quarter of all cetacean species are listed as threatened; *The IUCN Red List of Threatened Species*, 2024), reliable and efficient monitoring tools are urgently needed to support the conservation of cetacean diversity and their functional roles in marine ecosystems.

Cetacean distribution and occurrence is often determined through interviews with sea-faring communities and their traditional knowledge, visual and acoustic surveys, and, more recently, satellite image analysis (Bamford et al., 2020; Frasier et al., 2021; Liu et al., 2017). Visual monitoring methods are widely used to collect data suitable for various purposes such as species identification, abundance and distribution estimation, individual recognition, and estimation of body length (Boyd et al., 2019; Hammond et al., 2017; Thomas et al., 2010; Urian et al., 2015). However, such surveys are often expensive, rely on good weather windows, are labour intensive, and require the cetacean species under study at the surface frequently. Acoustic monitoring methods are also widely used as nearly all cetacean species produce a variety of vocalisations, often unique to that species and even to a population, for communication, navigation, and prey detection (Janik, 2014; Madsen et al., 2023). Acoustic data, like visual data, can be used to identify species, estimate population abundance and distribution, but if the species is not very vocal or its habitat is noisy, acoustic detection may be missed. Satellite imagery analysis is an emerging technique that has the potential to monitor large areas and remote habitats; however, at this time, standard analytical methodology is still being developed and the cost of purchasing satellite images or even tasking a satellite remains expensive (Bamford et al., 2020). Further, weather conditions must be good for clear images to be obtained (Höschle et al., 2021).

Given these challenges, eDNA analysis offers a new tool for detecting cetaceans. While eDNA analysis does not cover large spatial areas, as satellite imagery analysis can, it has several advantages over other methods: it can provide detailed genetic information, including intra-specific variation (Sigsgaard et al., 2017; Tsuji et al., 2023), requires relatively low on-site labor, and can detect cetaceans even when they are submerged or that have already left the immediate survey area. Additionally, eDNA concentration may serve as an estimate of biomass or abundance of target species (Rourke et al., 2022; Takahara et al., 2012), although careful study design and interpretation are essential (Rourke et al., 2022). Therefore, eDNA-based cetacean monitoring has the potential to enhance both the efficiency and effectiveness of cetacean monitoring.

Detecting cetaceans using eDNA analysis is a relatively new field. The widely used fish-targeting universal primer set, MiFish, which amplifies mitochondrial 12S rRNA gene (Miya et al., 2015), has been shown to detect cetacean eDNA (Closek et al., 2019). Other examples include the detection of cetacean eDNA using primers targeting the mitochondrial genome (Alter et al., 2022; Robinson et al., 2023; Zhang et al., 2023), such as the Riaz 12S primers (Riaz et al., 2011), MarVer primers (Valsecchi et al., 2020), and quantitative PCR approaches (Foote et al., 2012; Hashimoto et al., 2023) (For more examples, see Table 1 in Suarez-Bregua et al., 2022). Valsecchi et al. (2020) developed primer sets targeting marine mammals and other vertebrates, including cetaceans, pinnipeds, elasmobranchs, and bony fish. More recently, Deng et al. (2024) modified vertebrate universal primers and detected 18 cetacean species in the South China Sea. However, these primers were not specifically designed for cetaceans, leading to the amplification of more abundant non-cetacean eDNA, such as that from fish and common contaminants (e.g., human eDNA). This non-specific amplification can suppress the detection of target eDNA (Collins et al., 2019), and this issue is particularly important for cetacean eDNA metabarcoding because cetacean eDNA is typically present at low concentrations (e.g., Baker et al., 2018). These challenges underscore the need for novel primer sets that selectively amplify cetacean DNA while minimizing the amplification of other marine organisms, including fish, humans, and other vertebrates. Developing such primers is critical for establishing effective eDNA-based cetacean monitoring in marine ecosystems.

This study addresses the gap in cetacean-specific eDNA detection by developing and validating novel cetacean-targeting metabarcoding primer sets with minimal or no amplification of non-cetacean species. To achieve this, we collected mitochondrial genome sequences from cetaceans and related mammals from public databases, conducted *in silico* analyses to evaluate primer performance, and empirically tested multiple primer sets in both controlled and natural environments. Finally, we applied our newly developed primer sets to water samples collected from Hong Kong waters to detect the ‘vulnerable’ Indo-Pacific humpback dolphin (*Sousa chinensis*) (Jefferson & Smith, 2016). Our overarching aim is to establish a robust experimental protocol for cetacean eDNA monitoring, thereby contributing to the conservation of cetacean species.

## 2. Materials and Methods

### 2.1 Ethics statement

This study used cetacean tissue samples from stranded individuals found in coastal habitats or dead individuals in an aquarium to validate cetacean-specific primers for eDNA metabarcoding. All tissue samples were obtained with the necessary permits from the Okinawa Churashima Foundation (for all cetacean tissues except *Sousa chinensis*), the Agriculture, Fisheries and Conservation Department (AFCD) of Hong Kong, and Ocean Park Hong Kong (for *S. chinensis* samples). Seawater samples from dolphin pools were collected with permits from the Okinawa Churashima Foundation. No direct interaction with or sampling from live cetaceans occurred during this research.

### 2.2 Primer design

To design candidate primers, we retrieved mitochondrial genomes from 71 cetacean species, representing two suborders (Mysticeti and Odontoceti), 11 families, and 37 genera from the NCBI Reference Sequence Database (https://www.ncbi.nlm.nih.gov/refseq/; accessed 11 July 2023). Additionally, mitochondrial genomes of five mammal species (cattle [*Bos taurus*], wild boar [*Sus scrofa*], mouse [*Mus musculus*], cat [*Felis catus*], dog [*Canis lupus*], and human [*Homo sapiens*]) as well as other related taxa (dugong, manatee, and other marine mammals) were retrieved to assess potential amplification of non-target species (Table S1).

Retrieved sequences were aligned using the online version of MAFFT with default settings (Katoh & Standley, 2013; https://mafft.cbrc.jp/alignment/server/). Based on these alignments, 20 candidate regions were identified. Some primer sets were designed using `DesignPrimers()` function implemented in the DECIPHER package in R (Wright, 2016), while others were manually designed through visual inspection. In `DesignPrimers()` function, we tested several parameters such as `maxProductSize=500` and `numPrimerSets=30`. Suitable regions for cetacean eDNA metabarcoding were found in the 12S rRNA, 16S rRNA, and NAD3 genes (Fig. 1 and Table S2).

**Figure 1.**
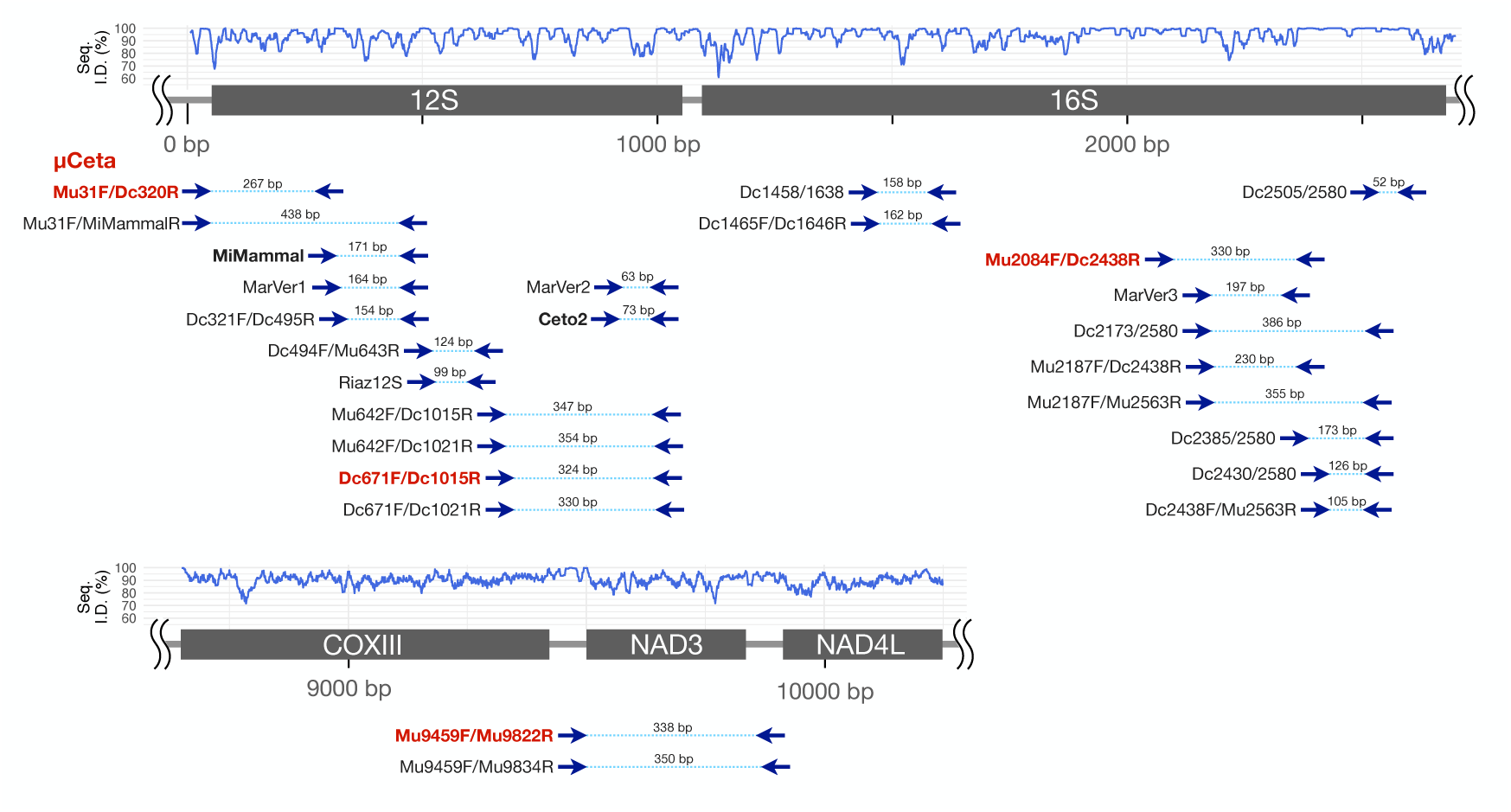
Candidate primers for Cetacea-specific eDNA metabarcoding designed from the mitochondrial genome. Previously reported mammal and vertebrate-targeting primers and our candidates primers for Cetacea-specific eDNA metabarcoding. “MiMammal,” “MarVer1,” “MarVer2,” “MarVer3,” “Ceto2” and “Riaz12S” are previously reported primers, while all other primers were designed in this study. Primers shown in boldface were empirically validated in this study, and newly designed primers among the empirically validated primers are highlighted in red. The prefixes “Mu” and “Dc” denote primers designed manually (by M. Ushio) and using DECIPHER, respectively. Each number between the primer regions indicates the target length in base pairs (bp) (i.e., the length of amplicon without primers). The “Seq. I.D. (%)” shown by the blue line indicates the 11-bp moving average of sequence identity (%). In the graph, “100%” means that all cetacean species have the same base at that position, while “70%” indicates that 70% of the cetacean species have the most common base. The base numbers are derived from the mitochondrial genome sequence of *Delphinus capensis*.

We designed the primer sets to amplify cetacean DNA while minimizing amplification of non-cetacean DNA (e.g., other marine mammals and common terrestrial species such as dogs and cats). In particular, we ensured that the primers do not amplify human DNA, as co-amplification of human DNA—a common contaminant, especially in coastal regions—could obscure the detection of low-concentration cetacean DNA. During primer design, we considered several technical best practices: primer length between 19 and 28 bp, GC% of 40– 60%, *T_m_* values of forward and reverse primers are as close as possible, and an amplicon length of less than 450 bp to achieve high sensitivity and compatibility with short-read sequencing.

### 2.3 *In silico* evaluation of the candidate primers

The performance of the candidate primers was evaluated *in silico* alongside other vertebrate-, mammal-, and cetacean-specific primers, including MiMammal (Ushio et al., 2017), MarVer1, MarVer2, MarVer3, Ceto2 (Valsecchi et al., 2020), and Riaz12S (Riaz et al., 2011). Ceto2 was included in our analysis as it was specifically designed as a cetacean primer set.

The characteristics and performance of the candidate primers were assessed using mitochondrial genomes of vertebrates retrieved from NCBI as of 19 July 2023. A total of 36,250 sequences from 10,855 vertebrate taxa were used for the *in silico* PCR which included 74 cetacean taxa (the same set of mitogenomes used for the primer design; as for the difference in the number of taxa, see “*The number of taxa in* in-silico *analysis*” for details), 2,378 non-cetacean mammal taxa, 1,556 bird taxa, 700 reptile taxa, 733 amphibian taxa, 5,411 fish taxa, and 1 additional taxon (*Branchiostoma belcheri*) (Table S3).

*In silico* PCR was conducted using “search_pcr” and “search_oligodb” commands implemented in USEARCH (Edgar, 2010). Specifically, we employed the following commands: `usearch -search_oligodb vertebrate_db.fa -db primer.fa -maxdiffs x -strand both -userout output.txt` (for each primer; `-maxdiffs` was examined up to 4) and `usearch -search_pcr vertebrate_db.fa -db primer_set.fa -maxdiffs x - strand both -minamp 30 -maxamp 2000 -pcrout output.txt -ampout ampout.txt`. (for each primer set; `-maxdiffs` was examined up to 3 for each primer set).

Based on the results of the *in silico* PCR results, we evaluated several key features of the primer sets: predicted amplicon lengths, the number of species and sequences potentially amplified, and the ability to differentiate closely related taxa. The predicted amplicon lengths and the number of amplified species and sequences were visualized using box and jitter plots. The ability to differentiate closely related taxa was assessed by calculating the edit distance between pairs of amplified species. The edit distance is a distance measure that quantifies the minimum number of single-character edits required to transform one DNA sequence into another. Therefore, if the edit distance between the amplicons of two different species was zero, the sequences were identical, making it impossible to differentiate between them.

### 2.4 Amplification of tissue-extracted DNA

Based on the *in silico* evaluation of the candidate primer sets (see Results), we selected four primer sets for empirical validation (Table 1). Mu31F/Dc320R and Dc671F/Dc1015R target the 12S rRNA gene, Mu2084F/Dc2438R targets the 16S rRNA gene, and Mu9459F/Mu9822R targets the NAD3 gene. Mu31F/Dc320R was later renamed “µCeta” (mάɪkroʊ-sɪtə) due to its superior performance in detecting cetacean species (see Results).

The amplification capacity of the four primer sets was initially tested using tissue-extracted DNA from 19 cetacean species belonging to 2 suborders and 5 families. Tissue (muscle) samples from all cetacean species, except the Indo-Pacific humpback dolphin (*S. chinensis*), were obtained from the Okinawa Churashima Foundation, where individual samples are preserved. Tissue samples of *S. chinensis* were archived by OPCFHK at Ocean Park Hong Kong. Both institutes collect and preserve stranded cetaceans for scientific research. DNA extraction protocols for the tissue samples and PCR conditions are described in the Supplementary Methods.

### 2.5 Water samples from aquarium and Hong Kong coastal regions

To evaluate the performance of the primer sets on water samples, we collected two types of water samples: Aquarium pool samples and Hong Kong coastal water samples. The first set of samples was collected from dolphin pools at the Okinawa Churaumi Aquarium using Sterivex filter cartridges (φ0.45-µm, SVHV010RS; Merck Millipore, Darmstadt, Germany) and the gravity filtration method (Oka et al., 2022). These water samples are hereafter referred to as “aquarium pool samples.” The aquarium has seven dolphin pools divided into two compartments, with water circulating within each compartment. The first compartment includes the “Main pool” (1,200 m^3^), “North Lagoon pool” (500 m^3^), and “South Lagoon pool” (500 m^3^). The second compartment consists of the “Main Show pool” (1,750 m^3^), “Medical Treatment pool” (500 m^3^), “Reproduction pool” (400 m^3^), and “Underwater Show pool” (500 m^3^). Five dolphin species, those are common bottlenose dolphin (*Tursiops truncatus*), Indo-Pacific bottlenose dolphin (*Tursiops aduncus*), false killer whale (*Pseudorca crassidens*), pygmy killer whale (*Feresa attenuate*), and rough-toothed dolphin (*Steno bredanensis*) were reared in the pools, and detailed information about the dolphin species is provided in Table S4.

From each pool, one or two 1 L seawater samples were collected (Table S4). Additionally, 1 L of MilliQ water was filtered as a field negative control. After filtration, 2 mL of RNAlater (Thermo Fisher Scientific, Waltham, MA, USA) was added, and samples were stored in a cooler box to preserve eDNA before transfer to a −20°C freezer in the laboratory. In total, we collected 10 seawater samples from the dolphin pools (nine pool samples and one field negative control).

Natural seawater samples were collected from Hong Kong coastal waters (approximately 22°10’N–22°30’N, 113°50’E–114°25’E). Three seawater samples, hereafter referred to as “preliminary Hong Kong water samples,” were taken from different locations to assess the amplification of target and non-target species. These samples were filtered using 0.45-µm Sterivex filter cartridges and plastic syringes: 500 ml of surface seawater at the first location (P1; 22°22’07”N, 114°25’55”E; The code “HK_P1” in Table S4) in February 2023 and 1 L of surface (“HK_P2_1”) and subsurface (“HK_P2_2”) seawater at the second location (P2; 22°20’27”N, 114°17’42”E) in July 2023. As a field negative control, 500-ml MilliQ water was filtered at the second location (“HK_nc”). After filtration, samples were stored in a cooler box to preserve eDNA before transfer to a −20°C freezer in the laboratory.

In addition to these preliminary seawater samples, we collected 500–1,500 mL of seawater samples in July 2024 from areas where Indo-Pacific humpback dolphins (*S. chinensis*) were sighted. These samples served as field positive controls, with two samples on 16 July 2024 (P3; 22°11’32”N, 113°50’57”E; “HK_P3_1” and “HK_P3_2”). The distance between *S. chinensis* individuals sighted and the sampling locations was approximately 30 m, leading us to expect that the seawater samples contained detectable amounts of *S. chinensis* eDNA. Hereafter, we refer to these as “natural positive samples.” To ensure contamination control, we also collected a MilliQ water sample as a field negative control. For all field samples, 2 ml of RNAlater was added and/or the samples were stored in a cooler box to preserve eDNA before transfer to a −20°C freezer in the laboratory.

### 2.6 eDNA extraction from seawater samples

Detailed eDNA extraction protocols are provided in the Supplementary Methods. Briefly, eDNA was extracted from Sterivex filter cartridges using a modified protocol based on a previous study (Fukuzawa et al., 2023). We used a DNeasy Blood & Tissue kit (Qiagen, Hilden, Germany), with the initial lysis step performed using Buffer ATL and Proteinase K. The extracted DNA was then purified following the manufacturer’s instructions and eluted in 100 µL of Buffer AE. To monitor contamination, at least one DNA extraction negative control (i.e., an empty filter cartridge) was included in each extraction process. Two replicate eDNA samples from the “Main Pool” and the “Main Show Pool” were pooled to generate a single eDNA sample for each pool, resulting in a total of seven aquarium pool eDNA samples. All eluted DNA samples were stored at −20°C until further processing.

### 2.7 Library preparation and sequencing

Detailed protocols for library preparation and sequencing are provided in the Supplementary Methods. For both aquarium pool samples and preliminary Hong Kong water samples, libraries were prepared using four candidate primer sets (Mu31F/Dc320R [µCeta], Dc671F/Dc1015R, Mu2084F/Dc2438R, and Mu9459F/Mu9822R).

The first-round PCR (1st PCR) was conducted to amplify the target regions of eDNA using Platinum SuperFi II PCR Master Mix (Thermo Fisher Scientific, Waltham, MA, USA) and forward and reverse primers with the Illumina sequencing primer (Table S5). Two PCR negative controls (using MilliQ water instead of template DNA) were included to monitor potential contamination during the library preparation process. After the 1st PCR, the PCR products for each sample were purified using AMPure XP (Beckman Coulter, Brea, CA, USA). The second-round PCR (2nd PCR) was then performed to append the Illumina adapters. Following the 2nd PCR, the indexed PCR products were pooled and the pooled product was purified. Then, the target-sized DNA of the purified library was excised, the double-stranded DNA concentration was adjusted, and the final library was sequenced on the Illumina NovaSeq platform (Illumina, San Diego, CA, USA) using a 2 × 250 PE reagent kit.

For natural positive samples, only the Mu31F/Dc320R (µCeta) primer set was used due to its superior performance (see Results and Discussion). In the library preparation process, we generally followed the same thermal cycler profile as that used for the aquarium samples. However, instead of using the 1st PCR primers with the Illumina sequencing primer, we employed “tagged” 1st PCR primers for more efficient library preparation (referred to as the “early-pooling protocol” in Ushio et al., 2022). The 1st PCR products were purified using ExoSAP-IT Express (Thermo Fisher Scientific, Waltham, MA, USA) and pooled for the 2nd PCR. The remaining processes including sequencing were identical to those described above.

In addition to the four candidate primer sets, we briefly tested the performance of previously developed primers, MiMammal (Ushio et al., 2017) and Ceto2 (Valsecchi et al., 2020), using aquarium and preliminary Hong Kong water samples. During library preparation, only the aquarium and preliminary Hong Kong water samples, along with PCR negative controls, were included. The library preparation method followed the standard 2-step PCR protocol, and the library was sequenced on the Illumina iSeq 100 system using iSeq 100 Reagent v2 (2 × 150 bp PE). For iSeq sequencing, 30% PhiX was spiked in to enhance sequencing quality.

In total, for testing the four primer sets, we had seven aquarium pool samples, one aquarium field negative control, three Hong Kong water samples (“HK_P1”, “HK_P2_1”, and “HK_P2_2” in Table S4), one Hong Kong water negative control sample, two DNA extraction negative controls, and two PCR negative controls (a total of 16 samples for each primer set). For testing the Mu31F/Dc320R (µCeta) primer set using natural positive samples, we had two Hong Kong water samples (“HK_P3_1” and “HK_P3_2” in Table S4), one Hong Kong water negative controls, one DNA extraction negative control, and two PCR negative controls (a total of six samples). For the MiMammal and Ceto2 primer sets, we had seven aquarium pool samples, three Hong Kong water samples, and two PCR negative controls (a total of 12 samples for each primer set).

### 2.8 Sequence data analysis

Detailed procedures for sequence data processing are available on Github (https://github.com/ong8181/micro-ceta). When the standard two-step PCR was used for library preparation, demultiplexing was performed using Illumina BaseSpace. When the early-pooling method was employed, demultiplexing was conducted using a custom script with cutadapt v4.7 (Martin, 2011).

The demultiplexed sequences were then quality-filtered using fastp v0.23.4 (Chen et al., 2018), and primer sequences were trimmed using cutadapt. The processed sequences were analyzed using DADA2 (Callahan et al., 2016), an amplicon sequence variant (ASV) approach implemented in R (R Core Team, 2023). During quality filtering, low-quality and unexpectedly short reads were removed. Error rates were estimated, and sequences were dereplicated, error-corrected, and merged to generate an ASV-sample matrix. Chimeric sequences were subsequently removed using the `removeBimeraDenovo` function in dada2 package. If multiple sequencing runs were conducted for the same library, `mergeSequenceTables` function was used after DADA2 processing to merge results. For the aquarium and preliminary Hong Kong water samples, we performed taxonomic identification using the ASV sequences, as these samples may contain eDNA from closely related species or hybrid species. For the natural positive samples, ASVs were clustered into operational taxonomic units (OTUs) at 97% similarity using the DECIPHER package in R (Wright, 2016).

Taxonomic identification of ASVs or OTUs was performed using two methods: the Query-centric auto-k-nearest-neighbor (QCauto) method (Tanabe & Toju, 2013) implemented in Claident v0.9.2021.10.22 (https://www.claident.org/) and `assignSpecies()` function implemented in the dada2 package, supplemented with a manual BLAST search (https://blast.ncbi.nlm.nih.gov/Blast.cgi). QCauto method is conservative; all nearest neighbors of a query sequence must share the same taxonomic information for taxa assignment. For example, if the QCauto method assigns a genus name to an ASV or OTU, all nearest neighbors surrounding the ASVs or OTUs must have the same genus name. This method increases the number of ASVs or OTUs without species names but reduces the likelihood of misassignments in species identification compared to other common methods (Sato et al., 2018; Q. Wang et al., 2007).

Given that we knew exactly which species were reared in the aquarium pools and that our target species included closely related species (e.g., *Tursiops truncatus*, *Tursiops aduncus*, and their hybrid *Tursiops truncatus* × *aduncus*), we also utilized the `assignSpecies()` function, using cetacean mitochondrial genomes retrieved from NCBI as a reference database. If the two approaches assigned different species names to an ASV or OTU, we adopted the result from the `assignSpecies()` function. In addition, sequences assigned to the genus “*Stenella*” were manually checked using BLAST, due to the presence of multiple closely related species. If species-level identification failed, sequences were labeled as “Unidentified Delphinidae” or “Unidentified Cetacea.” If an identified species was not reared in the aquarium pools, it was classified as “Non-present species.”

### 2.9 Statistical analysis

All statistical analyses were performed using R (R Core Team, 2023). For the aquarium pool and preliminary Hong Kong water samples, ASV matrices were rarefied using the coverage-based rarefaction (Chao & Jost, 2012) at a minimum coverage of 99.9%. The rarefied matrices were then converted to relative abundance (*i.e.*, percentage data), and rare ASVs (< 0.5%) were excluded from the analysis. Samples with fewer than 10 original sequences were excluded (i.e., most negative controls and some field samples). The relative abundance data were visualized using the ggplot2 package in R (Wickham, 2009). For the natural positive samples, only sequence reads were reported, as only one OTU was detected. All the code was deposited on Github (https://github.com/ong8181/micro-ceta).

## 3. Results and Discussion

### 3.1 *In silico* evaluation: Amplicon lengths

A total of 20 candidate primer sets were identified using the DECIPER package and manual inspection (Fig. 1 and Table S2). Most primer sets target the mitochondrial 12S rRNA and 16S rRNA gene regions, while two sets target the NAD3 gene region. The performance of the candidate primers was compared against other universal primers listed in Table S2.

Expected amplicon lengths for Cetacea (74 taxa), fish (5,411 taxa), and other vertebrates (5,368 taxa) were analyzed using *in silico* PCR, revealing median amplicon lengths ranging from approximately 100–500 bp (Fig. S1). All primer sets generated amplicons suitable for sequencing on short-read sequencers (e.g., Illumina). However, certain primer sets, such as Mu31F/MiMammalR, require 300 bp paired-end sequencing, which increases sequencing costs and can be performed with only a few Illumina sequencer (e.g., MiSeq and NextSeq 1000/2000).

### 3.2 *In silico* evaluation: primer mismatches in target/non-target taxa

All primer sets exhibited no more than one mismatch with the majority of cetacean DNA, indicating their strong amplification capability for most cetacean sequences (Fig. 2a; see individual primer results in Fig. S2).

**Figure 2.**
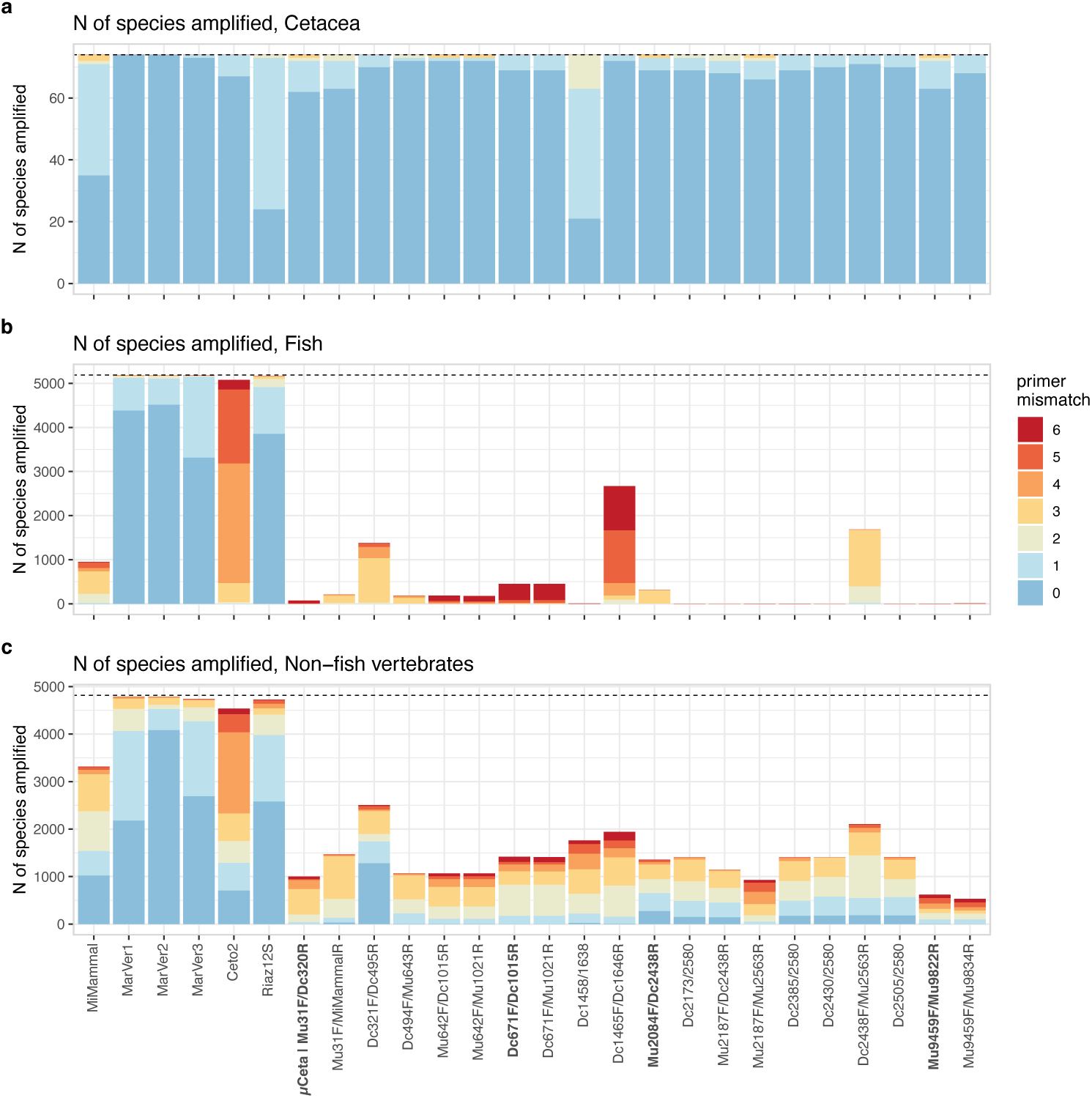
The number of species amplified by *in silico* PCR. The number of cetacean (**a**), fish (**b**), and non-fish vertebrate (**c**) species that can potentially be amplified using the previously reported and candidate primers. Colors represent the total number of mismatches between each primer set (forward and reverse primers) and the template DNA (more detailed results are provided in Figures S2—S4). The dashed horizontal line in each panel indicates the total number of species examined. Primer names shown in bold face are new primers that were empirically validated in this study.

The MarVer primes and Riaz12S showed few mismatches with fish and non-fish vertebrate DNA (Fig. 2b,c), consistent with their design as vertebrate-specific primer sets (Riaz et al., 2011; Valsecchi et al., 2020). The MiMammal (mammal-targeted primer set) and Ceto2 (cetacean-targeted primer set) exhibited three or more mismatches with most fish species, but could still amplify a significant number of non-fish vertebrates (Fig. 2b,c).

To further assess primer specificity, we analyzed the number of sequences amplified (Fig. S3) and the mismatch patterns between the forward/reverse primers and different taxa (Fig. S4). These analyses revealed similar patterns to those observed in Fig. 2. Importantly, the candidate primer sets we designed exhibited a greater number of mismatches with human DNA and common mammal DNA (e.g., dogs, cats, and others) as well as with fish, birds, reptiles, and amphibians compared to previously reported primers (Fig. S4). This suggests that our candidate primer sets have a lower likelihood of amplifying non-cetacean vertebrates. In this *in silico* evaluation, we regarded the non-amplification of human DNA as more important than that of other non-target DNA. Among the 20 candidate primer sets, 10 exhibited at least four mismatches with human DNA (i.e., Mu31F/Dc320R [µCeta], Mu642F/Dc1015R, Mu642F/Mu1021R, Dc671F/Dc1015R, Dc671F/Mu1021R, Mu2084F/Dc2438R, Mu2187F/Mu2563R, Dc2438F/Mu2563R, Mu9459F/Mu9822R, and Mu9459F/Mu9834R).

### 3.3 *In silico* evaluation: The capacity to differentiate cetacean species

The capacity of each primer set to differentiate cetacean species was evaluated by calculating the edit distance between their amplified products (Fig. 3). As expected, primer sets that amplify shorter regions, such as MarVer2, Ceto2, Riaz12S, Dc494F/Mu643R, and Dc2505/Dc2580, exhibited a reduced capacity to differentiate cetacean species. Specifically, more than 30–100 cetacean amplicon pairs generated by these primers shared identical sequences.

**Figure 3.**
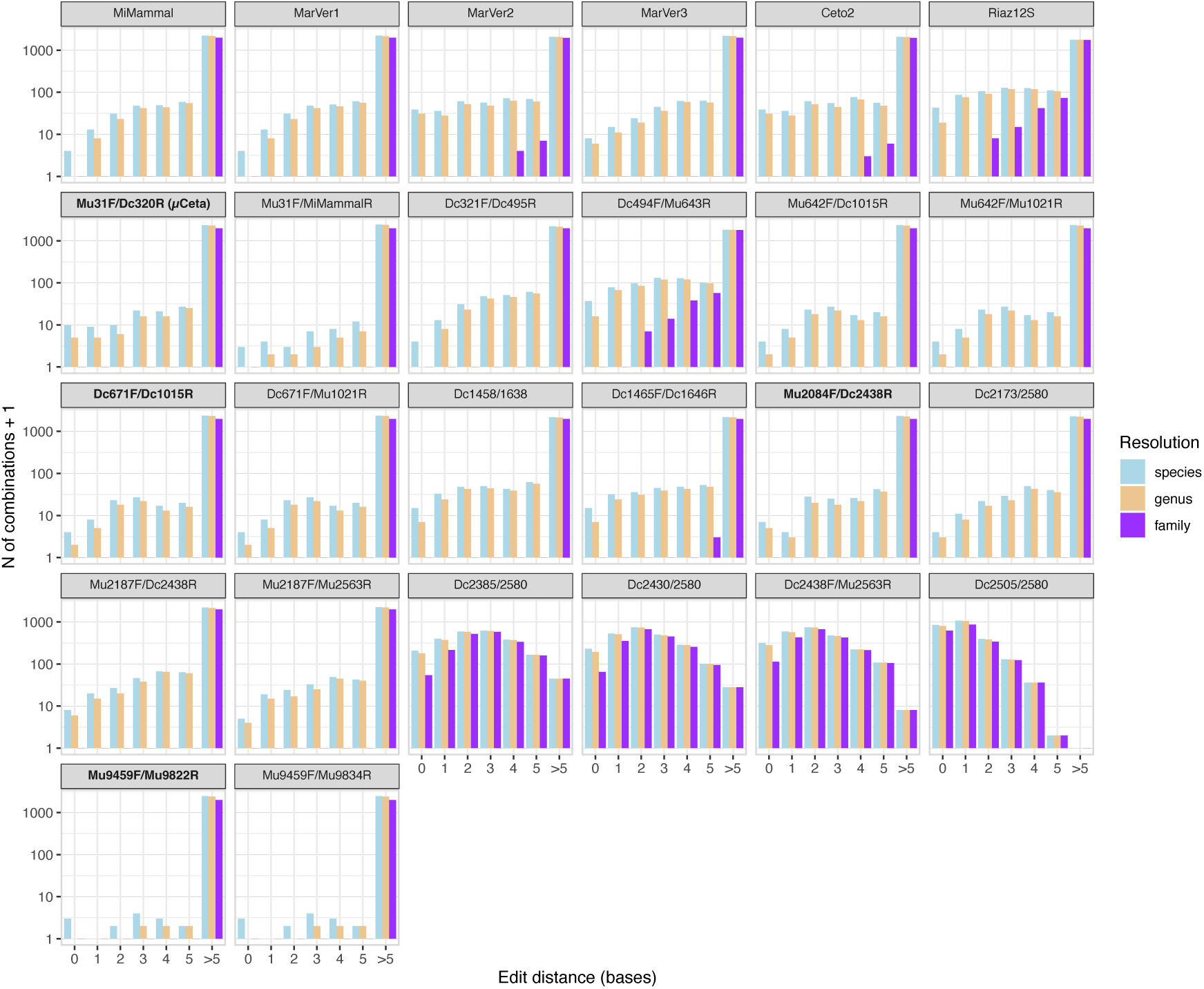
Capacity of previously reported and candidate primers to distinguish among different cetacean species, genera, or families. The capacity to distinguish different cetaceans was assessed by calculating the edit distances between amplicons through *in silico* PCR. The *x*-axis represents the edit distance, and the *y*-axis represents the log-transformed number of species pairs corresponding to each edit distance. Light blue bars indicate comparisons among different species, and light orange and purple bars represent comparisons among species belonging to different genera and families, respectively. The number of species pairs in the “0” edit distance category indicates the number of species pairs with identical sequences in the target region. Primer names shown in bold face are new primers that were empirically validated in this study.

Among the 10 primer sets with a low likelihood of amplifying human DNA, Dc2438F/Mu2563R exhibited a limited capacity to differentiate cetacean species, whereas the other primer sets resulted in fewer cetacean species pairs with identical sequences. From the nine primer sets that demonstrated high specificity for cetacean DNA and a superior capacity to differentiate cetacean species in the *in silico* analysis, we selected several primer sets for empirical validation. Preference was given to primers with fewer degenerate bases, as highly degenerate primers may exhibit lower amplification efficiency. The selected primers included: Mu31F/Dc320R (µCeta), Dc671F/Dc1015R, and Mu2084F/Dc2438R. Additionally, despite containing seven degenerate bases, we selected Mu9459F/Mu9822R for empirical validation due to its strong cetacean specificity and superior capacity to differentiate cetacean species. In total, we selected four primer sets for empirical validation (Table 1; see Fig. S5 for their primer logos).

### 3.4 Amplification of tissue-extracted cetacean DNA

We prepared tissue-extracted DNA from 19 cetacean species. The primer sets Mu31F/Dc320R (µCeta) and Mu2084F/Dc2438R successfully amplified all tissue-extracted DNA samples, while Dc671F/Dc1015R failed to amplify one species (*Globicephala macrorhynchus*), and Mu9459F/Mu9822R did not amplify four cetacean species (*Globicephala macrorhynchus*, *Kogia sima*, *Kogia breviceps*, and *Mesoplodon ginkgodens*) (Table 2). These results suggest that Mu9459F/Mu9822R may have lower amplification efficiency, potentially due to its higher number of degenerate bases. Another possible cause for the unsuccessful amplifications is DNA degradation, as the tissue samples of *G. macrorhynchus*, *K. sima*, and *M. ginkgodens* had been stored for 14, 12, and 15 years, respectively. Prolonged storage may have contributed to DNA degradation in combination with a higher number of degenerate bases in Mu9459F/Mu9822R, reducing amplification success.

### 3.5 eDNA metabarcoding for the aquarium pool and preliminary Hong Kong water samples

#### 3.5.1 Sequencing summary

We conducted eDNA metabarcoding on seawater samples collected from the aquarium pools and coastal regions of Hong Kong (Fig. 4). A total of 3,332,270 sequence reads were generated from 64 samples (16 libraries × four candidate primers; Table S6) by three NovaSeq runs, with an overall Q30 score of 92.08%. The four primer sets, Mu31F/Dc320R (µCeta), Dc671F/Dc1015R, Mu2084F/Dc2438R, and Mu9459F/Mu9822R produced 902,908 reads, 781,005 reads, 782,716 reads, and 865,741 reads, respectively. After quality filtering and DADA2 denoising, 2,572,852 reads remained (77.21% of the total reads). Overall, sequencing quality was sufficiently high (for more details, see Tables S6–S13).

**Figure 4.**
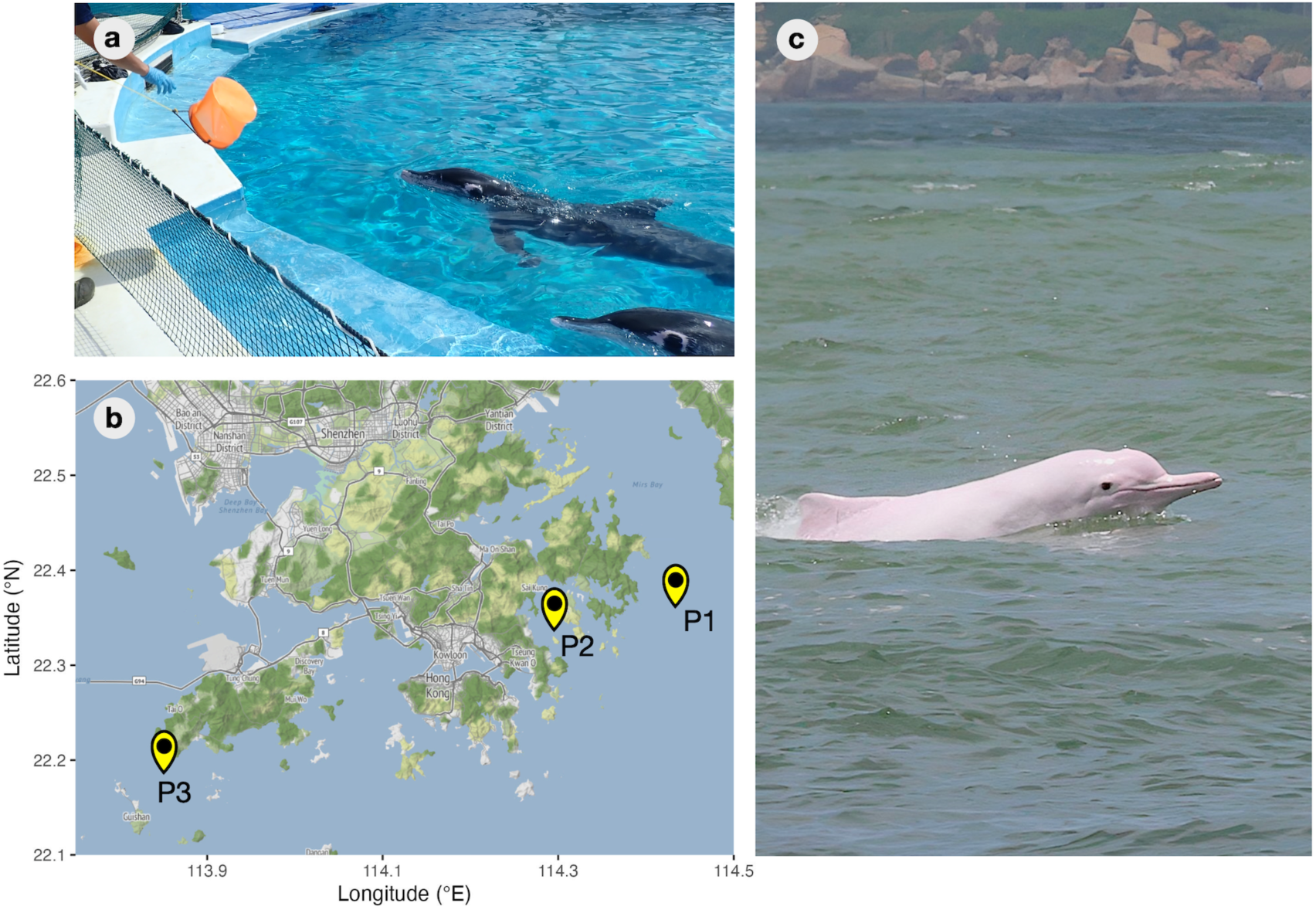
Collection of water samples at dolphin pools in Okinawa Churaumi Aquarium, field site locations in Hong Kong waters, and an Indo-Pacific Humpback Dolphin (*Sousa chinensis*) sighted during the field survey. (**a**) Water sampling at a dolphin pool in Okinawa Churaumi Aquarium for the “aquarium pool” experiment. (**b**) Water sampling locations in Hong Kong waters. P1 and P2 indicate the sampling locations for the preliminary testing, and P3 indicates the sampling locations for the positive field sample. Maps were created by ggmap package of R (©Stadia Maps, ©Stamen Design, ©OpenMapTiles and ©OpenStreetMap) (**c**) An Indo-Pacific Humpback Dolphins *(Sousa chinensis*) sighted during the field survey at site P3 (Photo by E. Matrai).

One of the highest read counts among negative controls was observed in DNA extraction negative controls amplified by Mu2084F/Dc2438R (2,650 reads; about 2–3% of the reads from other samples), and most of the sequence reads were derived from human DNA, which could be contaminated during the DNA extraction. Another negative control that generated some sequence reads was the field negative control collected from the Churaumi aquarium pools amplified by Mu31F/Dc320R (µCeta) (712 reads), but it accounted for at most 0.7% of the average reads from the pool samples. PCR negative controls had only a few reads (<0.1% of average sequence reads). These findings indicate that cross-contamination was not significant, and thus, no post-hoc decontamination analysis (e.g., using *decontam*; Davis et al., 2018) was performed.

#### 3.5.2 Detection of cetacean eDNA in the aquarium pools

First, we tested whether the four primer sets could detect cetacean species reared in the aquarium pools (Figs. 4a and 5; Table S4). In the Churaumi Aquarium pool test, *Pseudorca crassidens* and *Tursiops truncatus* (or *T. truncatus × aduncus*) were reared in four pools. All four primer sets successfully detected *P. crassidens* and *T. truncatus* in these pools (Fig. 5a). *Tursiops aduncus* was reared in three pools, where its eDNA was detected using the Mu31F/Dc320R (µCeta) and Mu9459F/Mu9822R primer sets. The majority of sequences identified as “Unidentified Delphinidae” in the North Lagoon and the Underwater Show Pool, detected by Dc671F/Dc1015R and Mu2084F/Dc2438R, were likely *T. aduncus*; however, the sequences could not be assigned to *T. aduncus* due to a lack of resolution in differentiating closely related species. eDNA from *Steno bredanensis* was detected by all candidate primer sets in the pool where *S. bredanensis* was reared. eDNA of *Feresa attenuata* was detected by all primer sets except Mu9459F/Mu9822R in the pool where *F. attenuata* was reared. Many sequences classified as “Unidentified Delphinidae” in the Underwater Show Pool detected by Mu9459F/Mu9822R were likely *F. attenuata*, yet these sequences could not be assigned to *F. attenuata*. These results suggest that all reared cetacean species were likely amplified by the four primer sets. However, only Mu31F/Dc320R (µCeta) was able to assign species names to all detected sequences.

**Figure 5.**
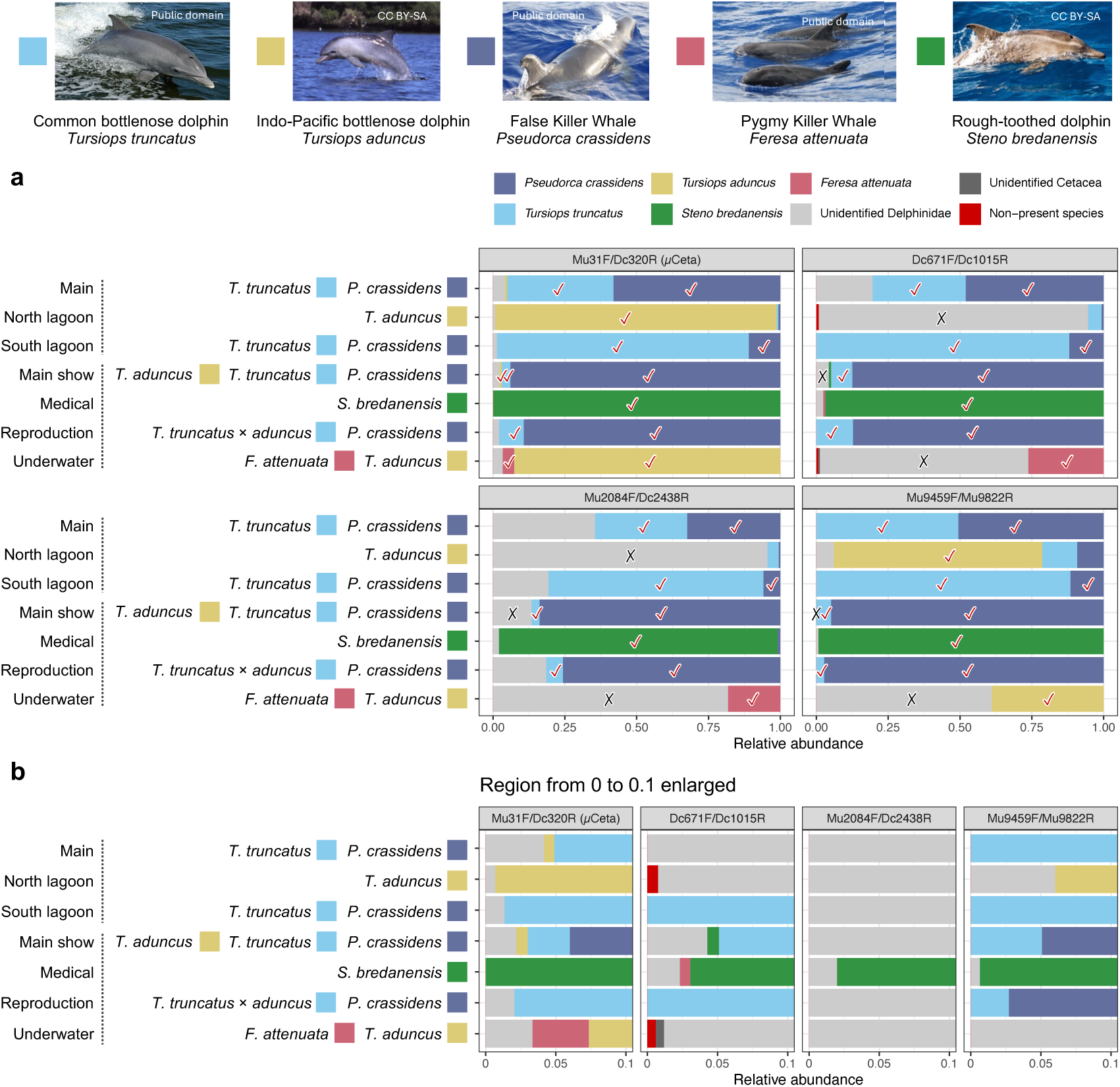
Cetacean eDNA metabarcoding results from the aquarium pool samples using the candidate primers. (**a**) Each panel indicates the results for four candidate primer sets. The *y*-axis represents seven dolphin pools from which seawater samples were collected, with species names and filled squares indicating the dolphin species reared in each pool. The *x*-axis represents the relative abundance of detected species eDNA. Colors represent different dolphin species or taxonomic categories when species identification is not possible. “Non-present species” denotes species detected that are not reared in the pool. A red tick (✓) indicates successful detection of target species. A black cross mark (×) indicates that, although the target species’ eDNA was seemingly amplified, its species name was not correctly assigned to the detected eDNA. Photo credits: *T. truncatus* (Public domain), *T. aduncus* (CC BY-SA 1.0, Aude Steiner), *P. crassidens* (Public domain), *F. attenuata* (Public domain), and *S. bredanensis* (CC BY-SA 3.0, Gustavo Pérez). All photos are sourced from Wikipedia. (**b**) The range from 0 to 0.1 is enlarged for clarity.

#### 3.5.3 Detection of non-present cetacean eDNA

We detected several cetacean species that were not present in each of the dolphin pools (see the enlarged figure in Fig. 5b). For instance, Mu31F/Dc320R (µCeta) detected eDNA of *T. aduncus* in the Main pool, and Dc671F/Dc1015R detected eDNA of *F. attenuata* in the Medical pool. The Main, North Lagoon, and South Lagoon pools belong to the same compartment, whereas the remaining pools belong to a separate compartment. Since water circulates within each compartment, cetacean eDNA may have been transported between pools. For example, *F. attenuata* eDNA could have moved from the Underwater Show pool to the Medical pool, resulting in detection via eDNA metabarcoding. Similar observations have been reported in other eDNA studies using aquariums (Kelly, Port, Yamahara, & Crowder, 2014; Miya et al., 2015). In some cases reported in these papers, the source of detection for non-present (referred to as “non-tank” in the papers) species was unknown; however, this underscores the sensitivity of eDNA metabarcoding and emphasizes the need for meticulous water sampling and library preparation to minimize cross-contamination.

The category of “Non-present species” includes *Balaenoptera acutorostrata*, which was detected only with Dc671F/Dc1015R, with a maximum proportion of 0.76%. This is negligible and does not impact our conclusions. Rare ASVs removed prior to this analysis included other non-present species such as *Tursiops australis* (*T. australis* is not recognized as a species, but here we follow the description of NC_022805 in the NCBI), *Globicephala macrorhynchus*, *Stenella longirostris*, and *Peponocephala electra*. The sources of these eDNA detections remain unknown, but some could be attributed to contamination from cetacean DNA attached to feeder fish.

#### 3.5.4 Supplemental eDNA metabarcoding: Detection of cetacean eDNA using MiMammal and Ceto2

To further assess species detection, we conducted eDNA metabarcoding on the Churaumi Aquarium pool samples and preliminary Hong Kong water samples using the MiMammal and Ceto2 primer sets (Fig. S6 and Table S6). A total of 2,406,398 sequence reads were generated from 24 samples (12 libraries using MiMammal and Ceto2 primers) in a single iSeq run, with an overall Q30 score of 95.80%. MiMammal and Ceto2 generated 1,078,756 and 1,327,642 reads, respectively. After quality filtering and DADA2 denoising, 2,212,982 reads (91.96%) remained for MiMammal and Ceto2 (Tables S6, S11, and S12). Both primer sets detected several cetacean species reared in the pools. However, some species could not be assigned species names, indicating that the capacity of these primer sets had lower species resolution than our candidate primer sets. Ceto2 exhibited particularly low taxonomic resolution, assigning species names to only two species, likely due to its short target length (73 bp), consistent with our *in silico* analysis (Fig. 3).

#### 3.5.5 Detection of non-target eDNA

We analyzed the detection of non-cetacean species from the aquarium pool samples and preliminary Hong Kong water samples (HK_P1, HK_P2_1, and HK_P2_2) (Fig. 4b) using the four candidate primer sets as well as the MiMammal and Ceto2 primer sets (Fig. S7). Raw ASV tables are available in Tables S14–S19.

Mu31F/Dc320R (µCeta), Dc671F/Dc1015R, and Mu9459F/Mu9822R detected almost exclusively cetacean eDNA. Mu31F/Dc320R (µCeta) and Dc671F/Dc1015R detected eDNA of the Indo-Pacific finless porpoise (*Neophocaena phocaenoides*) in the P1 surface water sample (HK_P1), a resident cetacean in Hong Kong waters (Jefferson & Moore, 2020). MiMammal primer set detected finless porpoise eDNA in P1, though at a relatively low proportion (Fig. S7). Despite our efforts to minimize amplification of non-cetacean eDNA, the Mu2084F/Dc2438R primer set detected a substantial amount of eDNA from common mammals (e.g., cows, pigs, and humans).

The MiMammal primer set detected a substantial number of human eDNA reads in all Hong Kong water samples, as expected. The Ceto2 primer set detected a large proportion of non-cetacean eDNA, including bacteria, non-cetacean mammals, and other eukaryotes (Fig. S7). The Ceto2 primer set was originally developed as a cetacean-specific primer set. However, its empirical validation was not conducted in Valsecchi et al. (2020), as it was not the primary objective of the study. Our empirical test suggests that Ceto2 may not be suitable for cetacean-targeting eDNA metabarcoding.

#### 3.5.6 Selection of µCeta (Mu31F/Dc320R) as the optimal primer set

Our empirical validation using the aquarium pool and preliminary Hong Kong water samples demonstrated that Mu31F/Dc320R effectively amplifies and differentiates most cetacean species tested in the present study (Fig. 5a) while exhibiting a low likelihood of amplifying non-cetacean species. Consequently, we designated Mu31F-Dc320R as µCeta (pronounced as mάɪkroʊ-sɪtə) and selected it for analyzing seawater samples collected from Hong Kong coastal areas where the Indo-Pacific humpback dolphin (*S. chinensis*) was sighted.

### 3.6 Detection of *Sousa chinensis* in Hong Kong waters

Two seawater samples were collected within approximately 30 m from where *S. chinensis* were sighted (P3 in Fig. 4c; HK_P3_1 and HK_P3_2 in Table S4). We then applied the µCeta primer set to these samples. A total of 3,539,281 sequence reads with an overall Q30 score of 92.48% were generated across three NovaSeq runs (Table S6). After quality filtering and DADA2 denoising, 3,246,690 reads remained (91.73% of the total reads). The substantial number of reads originated from seawater samples, while negative control samples for DNA extraction, PCR, and for field surveys showed negligible read counts (0 or 1 read per negative control).

Only one OTU was detected from the two seawater samples collected on 16 July 2024, and it was assigned to *S. chinensis* (HK_P3_1 and HK_P3_2; Table S20). From the samples, 1,799,357 and 1,446,733 reads of *S. chinensis* eDNA were identified, demonstrating that detection of *S. chinensis* is indeed possible using our eDNA-based approach. Importantly, no non-cetacean eDNA was detected in any of the three samples, suggesting that µCeta eDNA metabarcoding is highly sensitive for detecting this cetacean in natural seawater samples. Unfortunately, quantitative analysis of the detection sensitivity of µCeta primers is beyond the scope of this study (e.g., how close we should be to detect *S. chinensis* eDNA from a water sample), which should be addressed in future research.

### 3.7 Future direction of cetacean eDNA metabarcoding

Our *in silico* and empirical tests demonstrated that µCeta primers are well-suited for detecting cetacean eDNA from seawater samples. One clear limitation of our study is that we tested the performance of µCeta primers only on several toothed cetacean species using limited number of samples (five dolphin species in the aquarium pools and one dolphin species in Hong Kong waters). Although the *in-silico* evaluation suggested that µCeta can amplify most cetacean eDNA, some cetacean species exhibit multiple mismatches with µCeta primers (Table S21). For example, the Amazon River dolphin (*Inia geoffrensis*), bowhead whale (*Balaena mysticetus*), and 10 other cetacean species exhibit three, two, and one mismatches with µCeta primers, respectively. While the DNA of species with these numbers of mismatches could potentially be amplified with µCeta primers, empirical validation is necessary to ensure comprehensive detection of cetacean eDNA. Notably, human DNA exhibits only four mismatches with µCeta primers, yet we detected only two reads in the empirical testing, suggesting that even a few mismatches with µCeta primers might result in no amplification of target eDNA. Therefore, future research should validate the effectiveness of µCeta across a wider range of cetacean species and a broader array of field samples, including freshwater, deep-sea, and sediment samples.

Our results suggest that µCeta primers would detect cetacean eDNA from individuals located at least within tens of meters in Hong Kong waters. Passive acoustic monitoring has suggested that *S. chinensis* and *N. phocaenoides* inhabiting Hong Kong waters may be more active at night, when visual survey methodology is inappropriate (Wang & Hung, 2022). Consequently, µCeta metabarcoding could serve as an alternative method to study cetacean occurrence during periods of low visibility.

The abundance of cetacean species is critical information for conservation; however, this cannot be directly obtained from traditional eDNA metabarcoding. Species-specific qPCR can quantify cetacean eDNA (Hashimoto et al., 2023). Additionally, eDNA metabarcoding using spike-in DNA (Ushio et al., 2018) can provide an estimate of eDNA concentrations, serving as a rough index of cetacean abundance. Alternatively, eDNA metabarcoding with long-read sequencing could facilitate individual identifications based on intra-specific variation (Urban et al., 2023), which could enable direct estimation of the number of individuals under natural conditions.

Lastly, given that cetacean eDNA concentrations are expected to be lower than those of other marine vertebrates, such as fish, multiplexing µCeta primers with fish- or vertebrate-specific primers (Miya et al., 2015; Riaz et al., 2011; Valsecchi et al., 2020), or even developing new primers targeting other marine mammals, such as sirenians and pinnipeds, and multiplexing them, would enhance the efficiency of marine vertebrate monitoring using eDNA metabarcoding, while circumventing the adverse effects of amplification of non-target eDNA.

## 4. Conclusions

In this study, we developed cetacean-specific primers for eDNA metabarcoding and conducted *in silico* analysis to select four promising primer sets. These were empirically validated using aquarium pools and Hong Kong seawater samples, leading to the identification of µCeta—a primer set with high detection sensitivity, strong species differentiation capacity, and minimal non-target amplification.

The µCeta primer successfully detected the Indo-Pacific humpback dolphin (*S. chinensis*), a vulnerable species that resides within Hong Kong waters. Additionally, µCeta detected Indo-Pacific finless porpoise (*N. phocaenoides*), also classified as ‘vulnerable’ (*The IUCN Red List of Threatened Species*, 2024), in a seawater sample from the eastern Hong Kong waters. These findings highlight the potential of µCeta eDNA metabarcoding as a non-invasive and highly effective tool to monitor sparsely occurring and cryptic cetacean species.

Since µCeta was designed to detect a wide taxonomic range of cetacean species, its application can enhance cetacean monitoring efforts globally, supporting conservation initiatives and ecosystem management across various marine environments. Future studies should validate the effectiveness of µCeta using a broader range of field samples across diverse marine environments to further assess its applicability for global cetacean monitoring.

## Author contributions

MU conceived and designed research; MU and MM designed primers; MU, SO, SIO, TS, ROK, and MM collected tissues samples; MU, SO, and SIO collected aquarium water samples; MU, ROK, LP, and EM collected natural seawater samples; MU, SO, ROK, and TS performed experiments; MU analyzed data; MU and MM wrote the draft; All authors discussed the results and completed the manuscript.

## Data and code availability statements

All scripts used in the present study are available on Github (https://github.com/ong8181/micro-ceta) and Zenodo (https://doi.org/10.5281/zenodo.16139185). High-throughput sequence data was deposited in DDBJ Sequence Read Archives (DRA) (BioProject ID = PRJDB20491, DRA Run ID = DRR663608–DRR663784). Sanger sequence data was deposited in DDBJ Nucleotide Sequence Submission System (NSSS) (Accession No = LC863277–LC863344 and LC867741–LC867748).

## Acknowledgements

We thank Suixuan Huang, Ming-Wai Li, and Takamitsu Ohigashi for their assistance in the field work and Leung Kwan Chak for his assistance in the tissue sample collection and management. We thank Masayuki Sakata for his advice on the *in-silico* evaluation of the primer performance. We thank Haruna Okabe, Nozomi Kobayashi and the staff at the Okinawa Churashima Foundation, Okinawa Churaumi Aquarium, the Agriculture, Fisheries and Conservation Department (AFCD) of Hong Kong, and the Ocean Park Hong Kong for their permission for using cetacean tissue samples and assistance in collecting them. This research was supported by The Hong Kong University of Science and Technology Startup Fund to MU, and GRF16100724 from the Research Grants Council of the Hong Kong SAR, China to MU.

