## Supplementary Text for "µCeta: a set of cetacean-specific primers for environmental DNA metabarcoding with minimal amplification of non-target vertebrates"

#### Contents:

**Supplementary Methods.**

**Table S1.** List of mammal species for primer design.

**Table S2.** Primer sequences tested in this study.

**Table S3.** Queries to download vertebrate sequences for *in silico* PCR.

**Table S4**. List of aquarium dolphin pools and Hong Kong water samples for empirical validation of the primer sets.

**Table S5**. Primer sets for the Illumina library preparation.

**Table S6**. Sequence summary statistics.

**Table S7–S13**. Sequence processing summary for each primer set or field positive samples.

**Table S14–S19**. ASV-sample matrix for each primer set.

**Table S20**. OTU-sample matrix for field positive samples.

**Table S21**. The number of mismatches between Cetacea DNA and human DNA and µCeta primer set.

**Figure S1.** The distribution of expected amplicon lengths for each primer set.

**Figure S2.** The number of matched species for each primer by *in silico* PCR.

**Figure S3.** The number of amplified sequences by *in silico* PCR.

**Figure S4.** The relationships among the number of species, sequences, primer sets, taxa, and the number of mismatches examined by *in silico* PCR.

**Figure S5.** Primer logos for four candidate primers that were empirically validated.

**Figure S6.** Cetacean eDNA metabarcoding results from aquarium pool samples using MiMammal and Ceto2.

**Figure S7.** Detection of non-Cetacea environmental DNA using the four candidate and two previously reported primers.

### Supplementary Methods

##### *The number of taxa in* **in-silico** *analysis*

Counting the number of taxa used in the *in silico* analysis can be tricky, as the data we retrieved includes some synonyms and subspecies. For example, sperm whale has two species names in the database, *Physeter macrocephalus* and *Physeter catodon*. *Physeter catodon* was replaced with *Physeter macrocephalus* for the primer design. Also, Finless porpoise, *Neophocaena asiaeorientalis*, has three different taxa IDs, which were merged into one sequence for the primer design. In the *in-silico* PCR analysis, we counted the total number of taxa IDs, which was in total 10,855. For Cetacea, the total number of species used for the primer design was 71, while the total number of sequences used for the *in-silico* PCR was 74.

##### *DNA extraction for cetacean tissue samples*

DNA was extracted from cetacean tissues using a DNeasy Blood & Tissue kit (Qiagen, Hilden, Germany) following the manufacture's protocol. Briefly, cetacean tissues were cut into small pieces (up to 25 mg) and put into a 1.5-ml microtube. Then, Buffer ATL (180 µl) and Proteinase K (20 µl) were added to each tube. The tubes were incubated at 56˚C until the tissues were completely lysed (for 2–3 horus). The lysates were vortexed, and Buffer AL (200 µl) and absolute ethanol (200 µl) were added. The purification was performed following the manufacture's protocol. The purified DNA was eluted using 100 µl of Buffer AE and stored at –20˚C until further processing.

##### *Amplification of tissue-extracted DNA using the candidate primer sets*

For tissue-extracted DNA except Sousa chinensis, PCR was performed using Tks Gflex DNA polymerase (Takara, Otsu, Japan) following the manufacturer's protocol. The PCR was carried out with 30 cycles of a 7.7-µl reaction volume containing 0.2 µl Tks Gflex DNA polymerase, 3.8 µl PCR buffer, 0.4 µl of 5 µM F/R each primer (final concentration was 0.5 µM each), 2.3 µl sterile distilled H2O and 0.6 µl template. The thermal cycle profile after an initial 1 min denaturation at 94˚C was as follows: denaturation at 98◦C for 10 sec; annealing at 50˚C for 10 sec; and extension at 68˚C for 10 sec with the final extension at the same temperature for 7 min. The amplified products were purified using ExoSAP-IT (Thermo Fisher Scientific Baltics, UAB, Vilnius, Lithuania). Direct cycle sequencing was performed with dye-labelled terminators (BigDye terminator v. 1.1; Applied Biosystems, Foster City, CA, USA) following the manufacturer's protocol and the purified PCR products were sequenced for both strands on the ABI 3130xl Genetic Analyzer (Life Technologies, Carlsbad, CA, USA). The DNA sequences were edited and assembled using ATGC-MAC v.7.2.1 (Genetyx, Tokyo, Japan).

For tissue-extracted DNA of *Sousa chinensis*, PCR was performed using Platinum SuperFi II Master Mix (Thermo Fisher Scientific, Waltham, MA, USA) following manufacture's protocol. The PCR was carried out with a 20-µl reaction volume containing 10.0 µl of 2 × Platinum SuperFi II MasterMix, 2.0 µl of each 5 µM F/R primer (final concentration was 0.5 µM each), 4.0 µl of sterilized distilled $H_2O$, and 2.0 µl of template. The thermal cycle profile after an initial 30 sec denaturation at 98˚C was as follows (35 cycles): denaturation at 98˚C for 10 sec; annealing at 60˚C for 10 sec; and extension at 72˚C for 15 sec, with a final extension at the same temperature for 5 min. The amplified products were purified using ExoSAP-IT Express (Thermo Fisher Scientific, Waltham, MA, USA), and their concentrations were measured using Qubit dsDNA Quantification Assay Kits (Thermo Fisher Scientific, Waltham, MA, USA). The purified amplicons were sent to a sequencing company for Sanger Sequencing.

***DNA extraction from seawater samples***

eDNA was extracted from the Sterivex filter cartridges using a protocol described in a previous study (Fukuzawa et al., 2023) with some modifications. First, RNAlater solution was removed from the filter cartridge using a vacuum pump. One ml of $H_2O$ was added to wash the filter cartridge, which was removed using a vacuum pump. Then, Buffer ATL (380 µl) and Proteinase K solution (20 µl) were mixed, and the mixture was added to each filter cartridge. The materials on the cartridge filters were subjected to cell lysis by incubating the filters at 56˚C for 30 min. After the incubation, Buffer AL (400 µl) was added to the filter cartridge, and further incubated at 56˚C for 10 min to dissolve white precipitation. The incubated and lysed mixture was transferred into a new 2-ml tube from the of the filter cartridge using a manual centrifuge (Handzentrifuge, Hittich, Westphalia, Germany). The collected DNA was purified using a DNeasy Blood & Tissue kit following the manufacturer's protocol. After the purification, DNA was eluted using Buffer AE (100 µl). At least one DNA extraction negative control (i.e., an empty filter cartridge) was included in the DNA extraction process. Eluted DNA samples were stored at −20˚C until further processing.

***Detailed library preparation protocols***

For aquarium samples and natural seawater samples collected in the eastern area, libraries were prepared using the four candidate primer sets (Mu31F/Dc320R [µCeta] and Dc671F/Dc1015R, Mu2084F/Dc2438R, and Mu9459F/Mu9822R) and two previously developed primer sets (MiMammal and Ceto2). The PCR reagent compositions and thermal cycle profiles were the same for the normal 2-step protocol and the "early-pooling" protocol.

The 1st PCR was carried out with a 20-µl reaction volume containing 10.0 µl of 2 × Platinum SuperFi II PCR Master Mix (Thermo Fisher Scientific, Waltham, MA, USA), 2.0 µl of each 5.0 µM F/R primer with the Illumina sequencing primer (Table S5), 2.0 µl of MilliQ water and 4.0 µl of template. We used 4.0 µl of template DNA to increase the detection probability of low concentration eDNA (Ushio et al., 2022). Two PCR negative controls (i.e., we used MilliQ water instead of the template DNA) were included to monitor the potential contamination during the library preparation process. The thermal cycle profile after an initial 30 sec denaturation at 98˚C was as follows (35 cycles): denaturation at 98˚C for 10 sec; annealing at 60˚C for 10 sec; and extension at 72˚C for 15 sec, with a final extension at the same temperature for 5 min. The 1st-PCR products for each sample were purified using AMPure XP (PCR product: AMPure XP beads = 1:0.8) (Beckman Coulter, Brea, CA, USA).

Then, the 2nd PCR was carried out with a 20-µl reaction volume containing 10 µl of 2 × Platinum SuperFi II PCR Master Mix, 2.0 µl of each 5.0 µM index F/R primer (Table S5), and 2.0 µl of the purified 1st PCR product. The thermal cycle profile after an initial 30 sec denaturation at 98˚C was as follows (10 cycles): denaturation at 98˚C for 10 sec; annealing/extension at 72˚C for 15 sec, with a final extension at the same temperature for 5 min.

After the 2nd PCR, the indexed PCR products were combined. The pooled 2nd PCR product was purified using AMPure XP (PCR product: AMPure XP beads = 1:0.8;). The target-sized DNA of the purified library was excised using E-Gel SizeSelect (ThermoFisher Scientific, Waltham, MA, USA). The double-stranded DNA concentrations of the libraries were quantified using a Qubit dsDNA Quantification Assay Kit (Thermo Fisher Scientific, Waltham, MA, USA). The double-stranded DNA concentration of the combined library was then adjusted and sequenced on the Illumina NovaSeq platform (Illumina, San Diego, CA, USA) using 2 × 250 PE reagent kit or the Illumina iSeq 100 platform using 2 × 150 PE reagent kit.
