## Supplementary Figures for "µCeta: a set of cetacean-specific primers for environmental DNA metabarcoding with minimal amplification of non-target vertebrates"

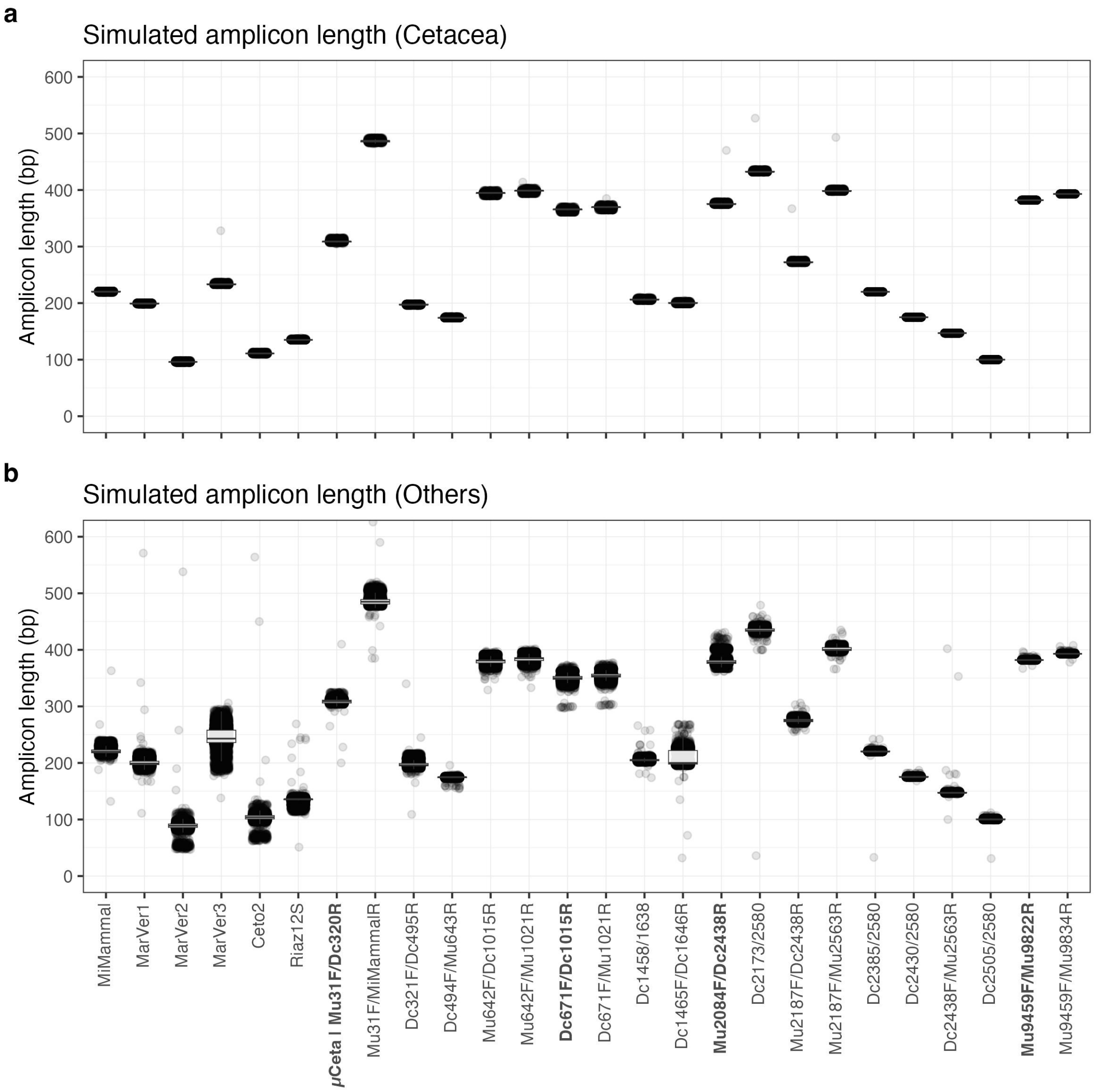


**Figure S1 | The distribution of expected amplicon lengths for each primer set.** Expected amplicon lengths for Cetacea species (**a**) and other species (**b**). Results are based on *in silico* PCR analysis of 71 cetacean species and 36,250 non-cetacean vertebrates. Each horizontal bar represents the median expected amplicon length for each primer set. Primer names shown in bold face are new primers that were empirically validated in this study.

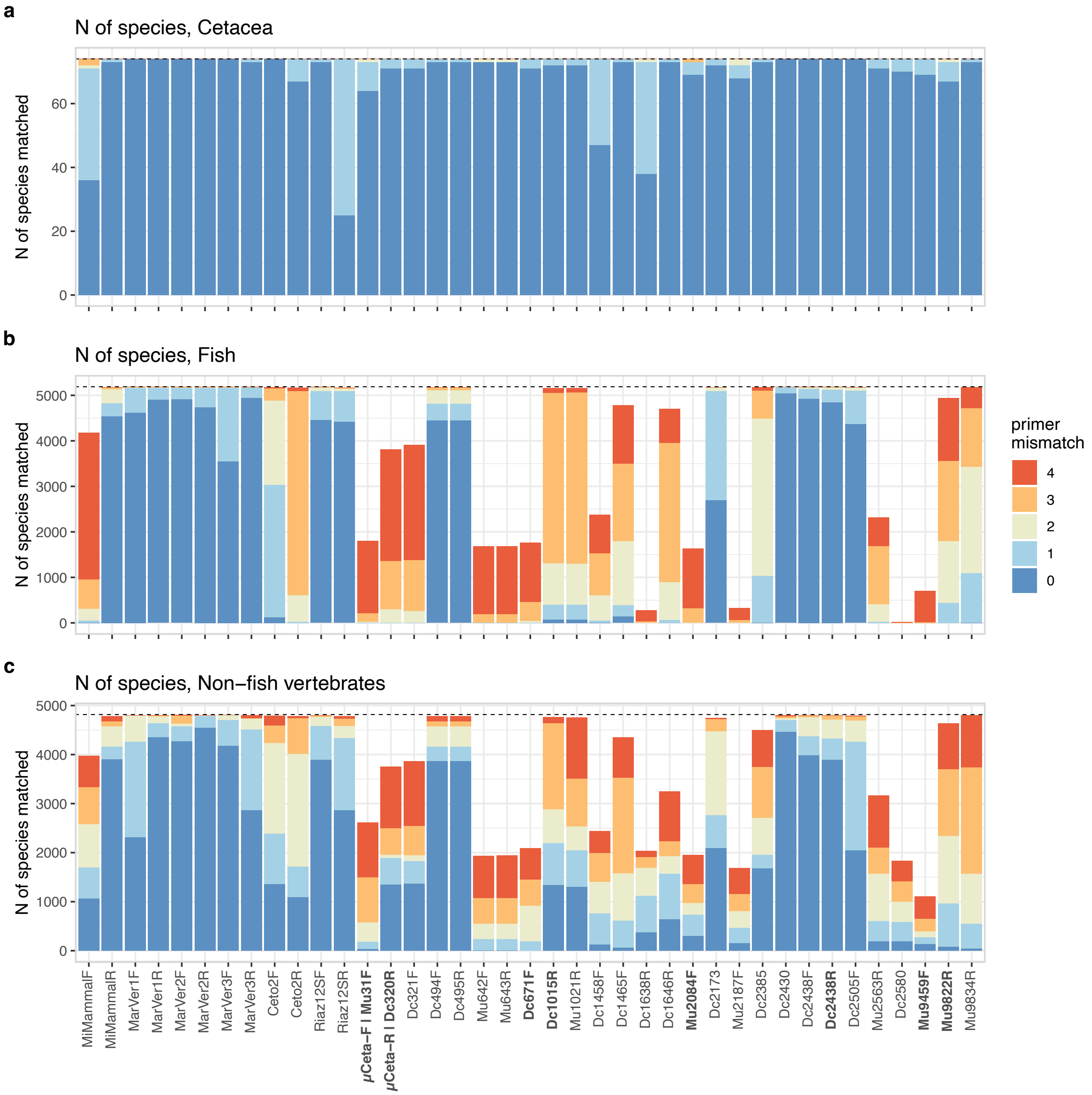


**Figure S2 | The number of matched species for each primer by *in silico* PCR.** The number of cetacean (**a**), fish (**b**), and non-fish vertebrate (**c**) sequences that are matched with each primer. Colors indicate the total number of mismatch between a primer set (forward and reverse primers) and template DNA. The dashed horizontal lines indicate the total number of sequences examined. Primer names shown in bold face are new primers that were empirically validated in this study.

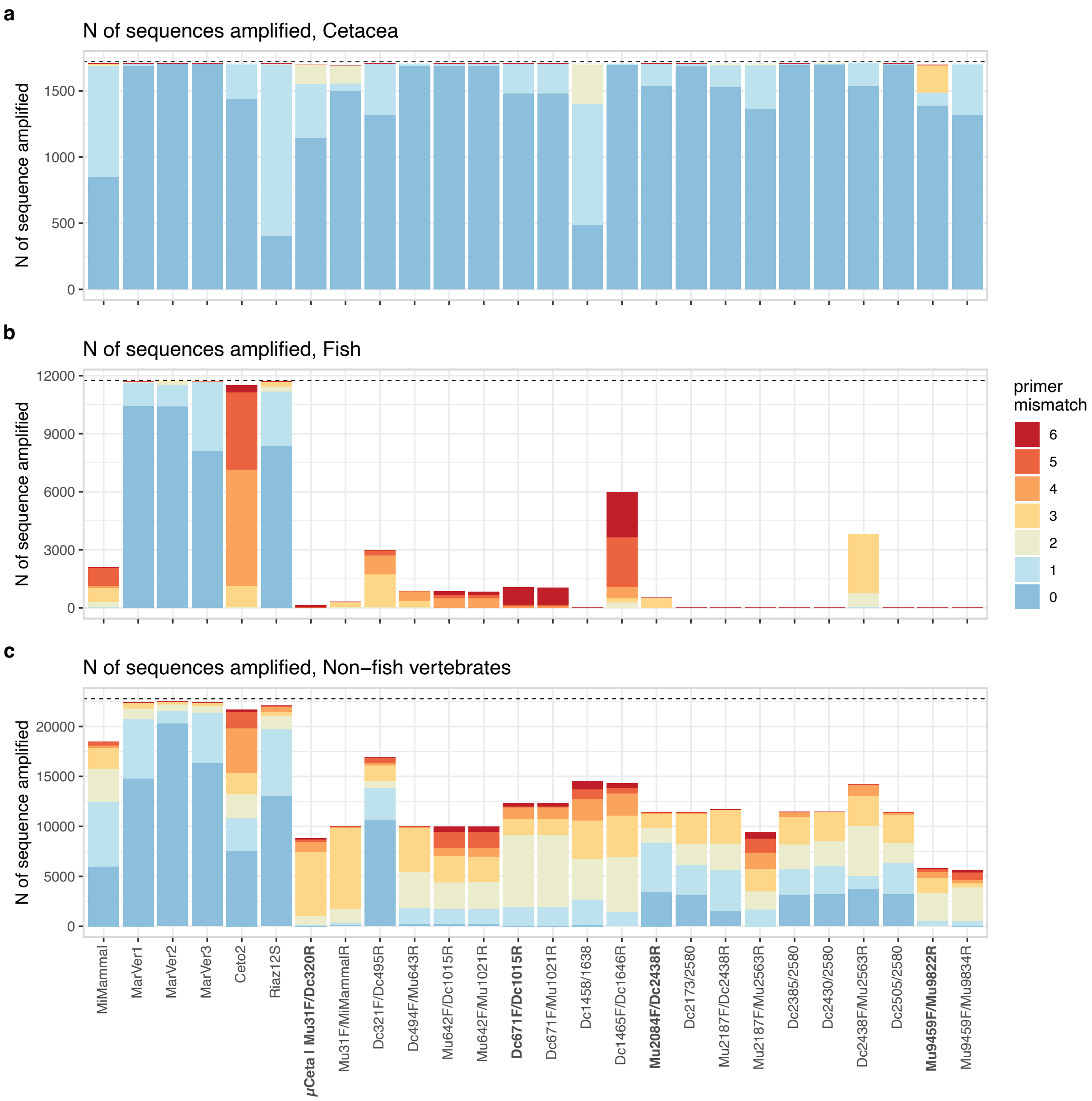


**Figure S3 | The number of amplified sequences by *in silico* PCR.** The number of cetacean (**a**), fish (**b**), and non-fish vertebrate (**c**) sequences potentially amplified by the previously reported and candidate primers. Colors indicate the total number of mismatches between the primer set and the template DNA (mismatches for forward and reverse primers were combined). The dashed horizontal lines represent the total number of sequences examined. Primer names shown in bold face are new primers that were empirically validated in this study.

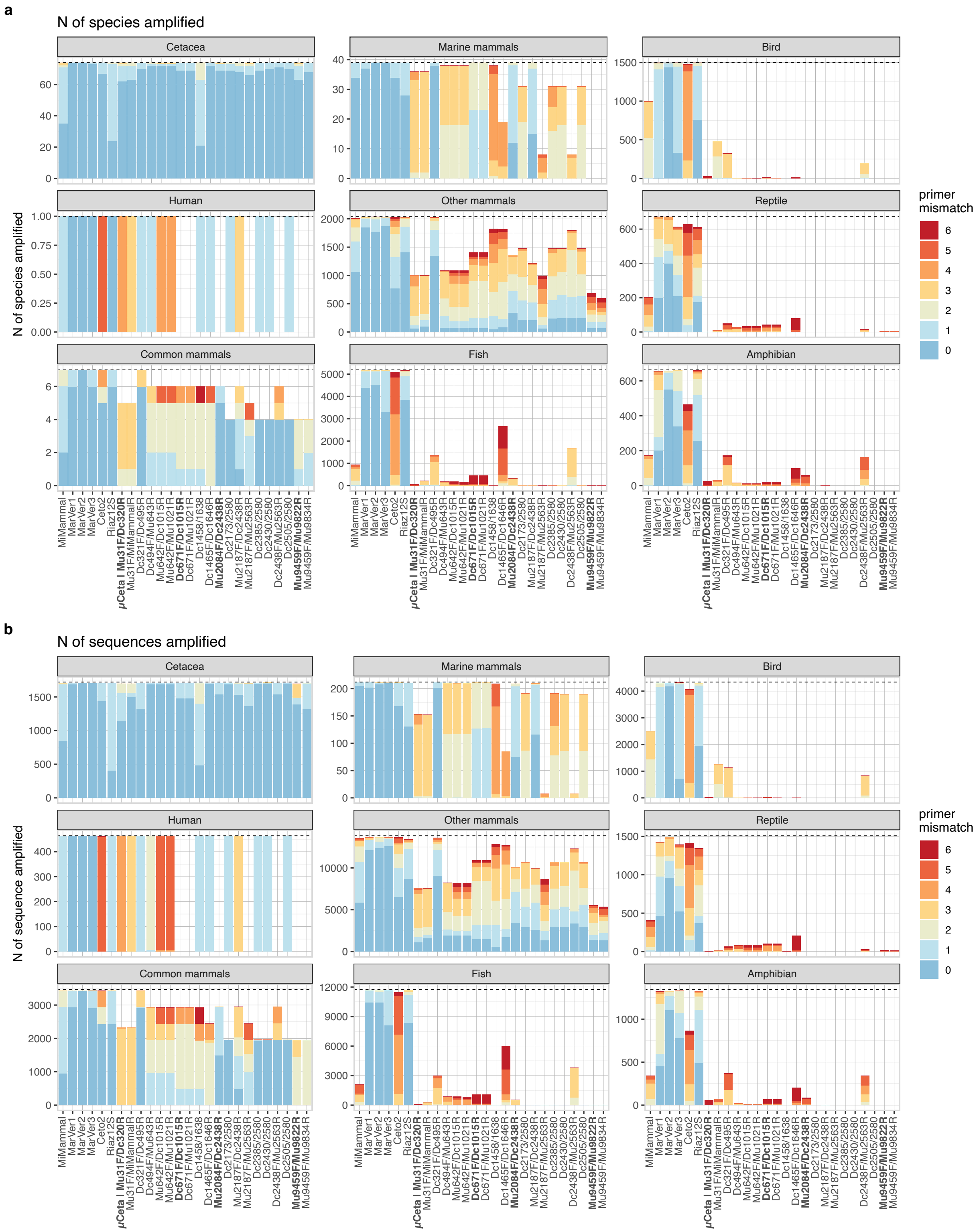


**Figure S4 | The relationships among the number of species, sequences, primer sets, taxa, and the number of mismatches examined by *in silico* PCR.** The number of mismatches between the template DNA and primers examined by *in silico PCR*, summarized by the number of species (**a**) and sequences (**b**). Each panel represents a different taxonomic group (denoted in the strip). Colors indicate the total number of mismatches. The dashed horizontal lines represent the total number of species examined. Primer names shown in bold face are new primers that were empirically validated in this study.

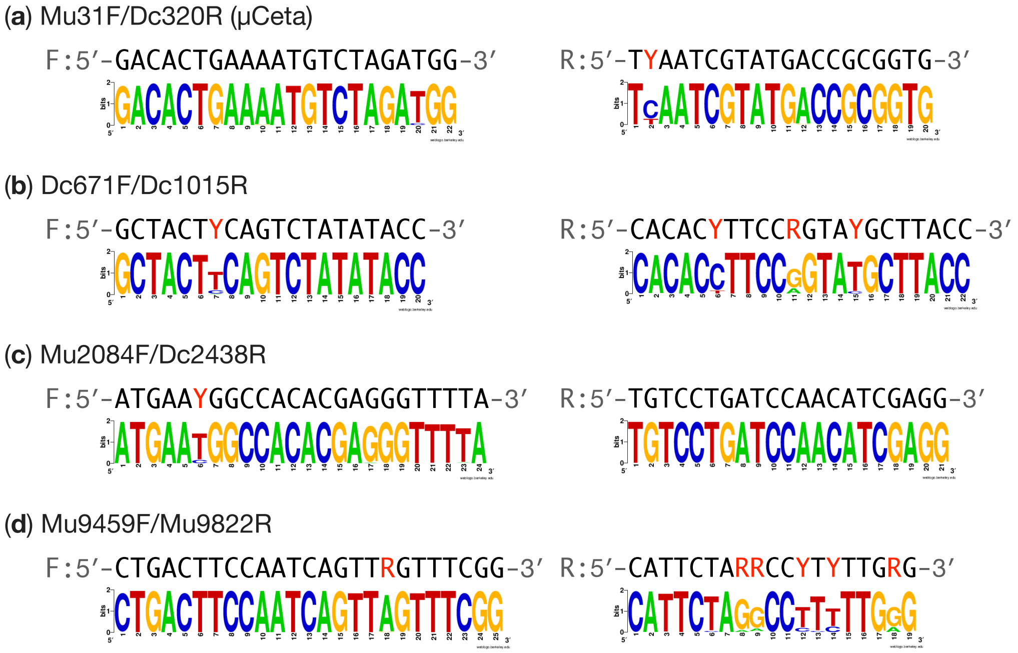


**Figure S5 | Primer logos for four candidate primers that were empirically validated.** Primer logos for (**a**) Mu31F/Dc320R ($\mu$Ceta), (**b**) Dc671F/Dc1015R, (**c**) Mu2084F/Dc2438R, and (**d**) Mu9459F/Mu9822R. Colors represent different bases, with red characters indicating degenerate bases. In each primer set, "F:" and "R:" denote the sequences for forward and reverse primers, respectively. The $y$-axis represents the information (in bits) regarding the conservation of the base at each position. A value of "2" indicates that all the bases are the same, while a value of "0" indicates that the bases are random.

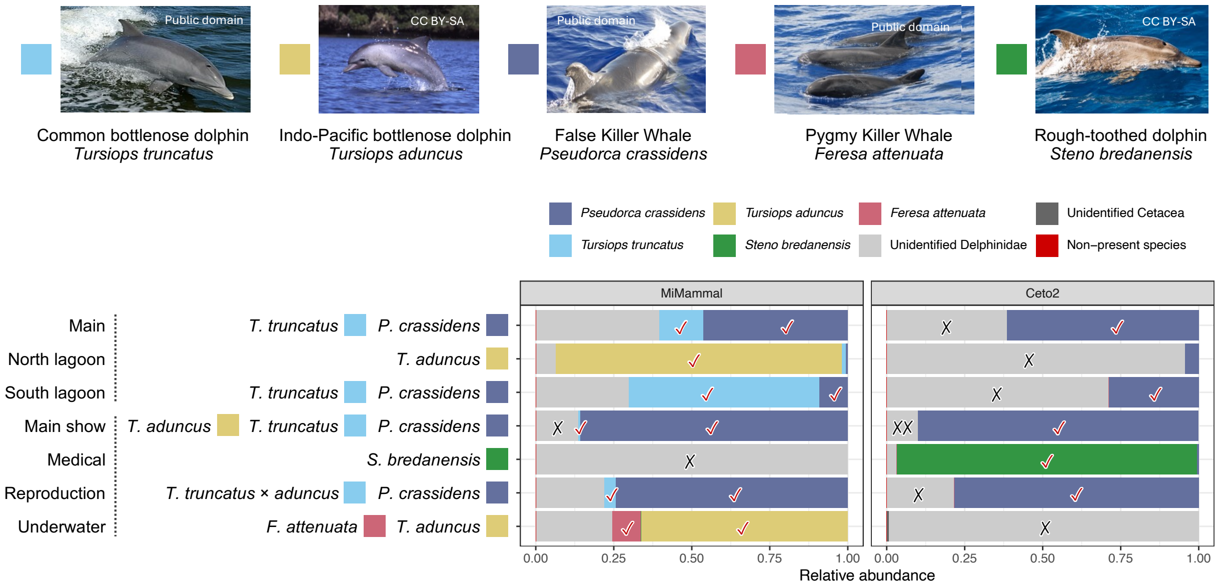


**Figure S6 | Cetacean eDNA metabarcoding results from aquarium pool samples using MiMammal and Ceto2.** Each panel presents the results of two previously reported primer sets, MiMammal and Ceto2. The $y$-axis represents seven dolphin pools from which seawater samples were collected, with species names and filled square indicating the dolphin species reared in each pool. The $x$-axis represents the relative abundance of detected species eDNA. Colors represent different dolphin species or taxonomic categories when species identification is not possible. "Non-present species" denotes species detected that are not reared in the pool. A red tick (✓) indicates successful detection of target species. A black cross mark (×) indicates that, although the target species' eDNA was seemingly amplified, its species name was not correctly assigned to the detected eDNA. Photo credits: *T. truncatus* (Public domain), *T. aduncus* (CC BY-SA 1.0, Aude Steiner), *P. crassidens* (Public domain), *F. attenuata* (Public domain), and *S. bredanensis* (CC BY-SA 3.0, Gustavo Pérez). All photos are sourced from Wikipedia.

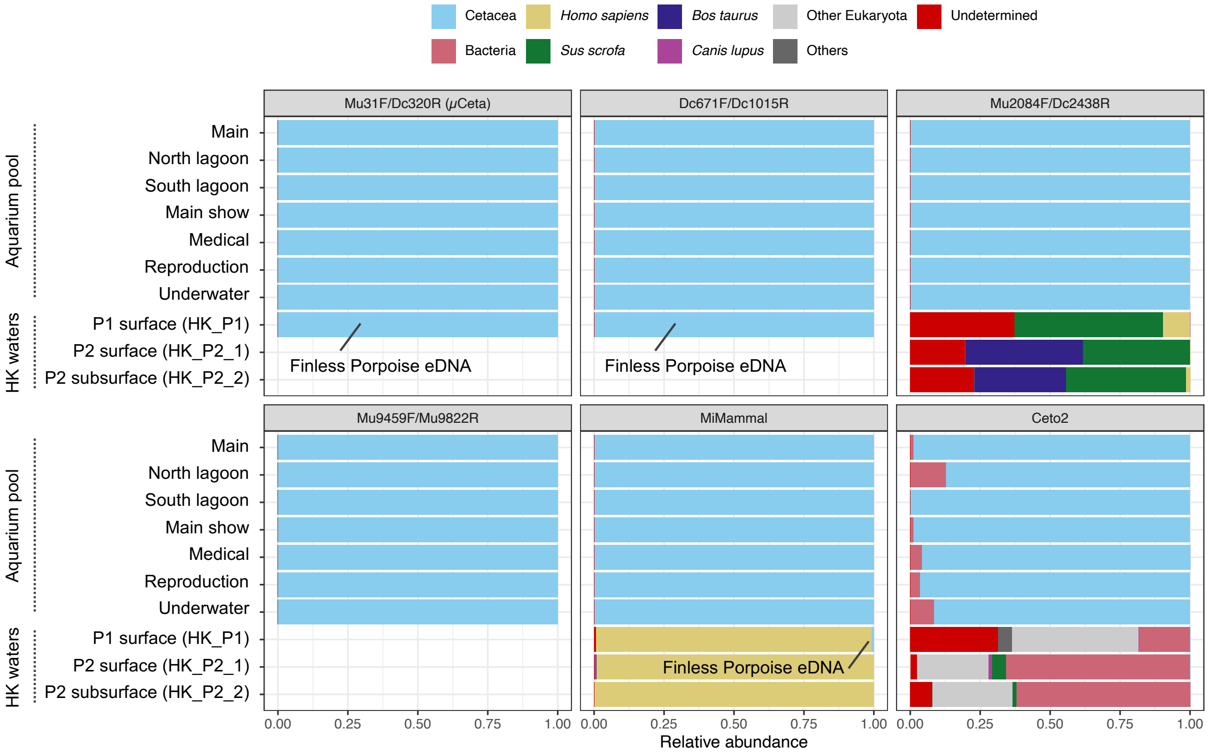


**Figure S7 | Detection of non-Cetacea environmental DNA using the four candidate and two previously reported primers.** Each panel presents the results for four candidate and two previously reported primer sets. The $y$-axis represents seven dolphin pools and three natural seawater samples. The $x$-axis represents the relative abundance of detected species eDNA. Colors represent taxonomic categories.
